# Multi-rate Poisson Tree Processes for single-locus species delimitation under Maximum Likelihood and Markov Chain Monte Carlo

**DOI:** 10.1101/063875

**Authors:** P. Kapli, S. Lutteropp, J. Zhang, K. Kobert, P. Pavlidis, A. Stamatakis, T. Flouri

**Affiliations:** The Exelixis Lab, Scientific Computing Group, Heidelberg Institute for Theoretical Studies, Schloss-Wolfsbrunnenweg 35, D-68159 Heidelberg, Germany; Department of Informatics, Institute of Theoretical Informatics, Karlsruhe Institute of Technology, 76128 Karlsruhe, Germany; Institute of Molecular Biology and Biotechnology (IMBB), Foundation of Research and Technology, Nikolaou Plastira 100 GR-70013 Heraklion, Crete, Greece 76128 Karlsruhe, Germany

## Abstract

**Motivation:** In recent years, molecular species delimitation has become a routine approach for quantifying and classifying biodiversity. Barcoding methods are of particular importance in large-scale surveys as they promote fast species discovery and biodiversity estimates. Among those, distance-based methods are the most common choice as they scale well with large datasets; however, they are sensitive to similarity threshold parameters and they ignore evolutionary relationships. The recently introduced “Poisson Tree Processes” (PTP) method is a phylogeny-aware approach that does not rely on such thresholds. Yet, two weaknesses of PTP impact its accuracy and practicality when applied to large datasets; it does not account for divergent intraspecific variation and is slow for a large number of sequences.

**Results:** We introduce the multi-rate PTP (mPTP), an improved method that alleviates the theoretical and technical shortcomings of PTP. It incorporates different levels of intraspecific genetic diversity deriving from differences in either the evolutionary history or sampling of each species. Results on empirical data suggest that mPTP is superior to PTP and popular distance-based methods as it, consistently, yields more accurate delimitations with respect to the taxonomy (i.e., identifies more taxonomic species, infers species numbers closer to the taxonomy). Moreover, mPTP does not require any similarity threshold as input. The novel dynamic programming algorithm attains a speedup of at least five orders of magnitude compared to PTP, allowing it to delimit species in large (meta-) barcoding data. In addition, Markov Chain Monte Carlo sampling provides a comprehensive evaluation of the inferred delimitation in just a few seconds for millions of steps, independently of tree size.

**Availability:** mPTP is implemented in C and is available for download at http://github.com/Pas-Kapli/mptp under the GNU Affero 3 license. A web-service is available at http://mptp.h-its.org.

## 1 INTRODUCTION

Species are fundamental units of life and form the most common basis of comparison in evolutionary studies. Therefore, species delimitation is a critical task in systematic studies with potential implications in all subfields of biology that involve evolutionary relationships. In line with the species concept of De Queiroz (2007) and the integrative taxonomy approach (Dayrat, 2005), a reliable identification of a new species involves data from multiple sources (e.g., ecology, morphology, evolutionary history). Such an approach is necessary for meticulous comparative evolutionary studies but is tremendously difficult to apply, if at all possible, in large biodiversity studies (e.g., vast barcoding, environmental, microbial samples). In such studies, researchers use alternative units of comparison that are easier to delimit [Molecular Operational Taxonomic Units (Blaxter *et al.*, 2005), Recognizable Taxonomic Units (Oliver and Beattie, 1993)] rather than going through the cumbersome task of delimiting and describing full species. The correspondence of these units to real species remains ambiguous (Casiraghi *et al.*, 2010), especially in the absence of additional information. However, they are practical for biodiversity and *β*-diversity estimates (Valentini *et al.*, 2009).

With the introduction of DNA-barcoding (Hebert *et al.*, 2003) and the advances in coalescent theory (Hickerson *et al.*, 2010), genetic data became the most popular data source for delimiting species. Several algorithms and implementations for molecular species delimitation exist, most of which are inspired by the phylogenetic species concept (Zhang *et al.*, 2013; Fujisawa and Barraclough, 2013; Yang and Rannala, 2014) and the DNA barcoding concept (Puillandre *et al.*, 2012; Hao *et al.*, 2011; Edgar, 2010). These methods address different research questions (Casiraghi *et al.*, 2010); thus, the user needs to assess several factors before choosing the most appropriate method for a particular study. The “species-tree” approaches rely on multiple genetic loci (Yang and Rannala, 2014; Jones *et al.*, 2015) and account for potential species-tree/gene-tree incongruence (Maddison, 1997). Such methods are the most appropriate when the goal is to identify and describe new species. However, most current implementations (Yang and Rannala, 2014; Jones *et al.*, 2015) are computationally demanding and can only be applied to small datasets of closely related taxa, becoming impractical with a growing number of species and/or loci [see Fujisawa *et al.* (2016) for a recently introduced faster method]. Large-scale biodiversity (meta-) barcoding studies comprise hundreds or even thousands of samples of high evolutionary divergence. The goal of such studies often is to obtain *β*-diversity estimates (comparative studies ofdifferent treatments, ecological factors, etc.) or a rough estimate of the biodiversity for a given sample. Hence, a researcher may use distance-based methods (Puillandre *et al.*, 2012; Hao *et al.*, 2011; Edgar, 2010) that scale well on large datasets with respect to run times. However, these methods ignore the evolutionary relationships of the involved taxa and rely on not necessarily biologically meaningful *ad hoc* sequence similarity thresholds. Moreover, they are restricted to single-locus data, which reduces species delimitation accuracy (Dupuis *et al.*, 2012).

The General Mixed Yule Coalescent (GMYC; Pons *et al.*, 2006; Fujisawa and Barraclough, 2013) and the recently introduced Poisson Tree Processes (PTP; Zhang *et al.*, 2013) are two similar models that bridge the gap between “species-tree” and distance-based methods for species delimitation. While GMYC and PTP are also restricted to single loci, they do take the evolutionary relationships of the sequences into account. Since, at the same time, they are computationally inexpensive they can be deployed for analyzing large (meta-)barcoding samples. The GMYC method (Fujisawa and Barraclough, 2013) uses a speciation (Yule, 1925) and a neutral coalescent model (Hudson, 1990). It strives to maximize the likelihood score by separating/classifying the branches of an ultrametric tree (in units of absolute or relative ages) into two processes; within and between species. In contrast to GMYC, PTP models the branching processes based on the number of accumulated expected substitutions between subsequent speciation events. PTP tries to determine the transition point from a between-to a within-species process by assuming that a two parameter model, with one parameter for speciation and one parameter for the coalescent process best fits the data. The underlying assumption is that each substitution has a small probability of generating a branching event. Within species, branching events will be frequent whereas among species they will be more sparse.The probability of observing *n* speciations for *k* substitutions follows a Poisson process and therefore, the number of substitutions until the next speciation event can be modeled via an exponential distribution (Zhang *et al.*, 2013). Given that PTP directly uses substitutions, it does not require an ultrametric input tree, the inference of which can be time consuming and error prone. Thus, PTP often yields more accurate delimitations than GMYC (Tang *et al.*, 2014).

Here, we introduce a new algorithm and an improved model as well as implementation of PTP that alleviates previous shortcomings of the method. The initial PTP assumes only one exponential distribution for the speciation events and only one for the coalescent events, across all species in the phylogeny. While the speciation rate can be assumed to be constant among closely related species, the intraspecific coalescent rate and consequently the genetic diversity may vary significantly even among sister species. This divergence in intraspecific variation can be attributed to factors, such as population size and structure, population bottlenecks, selection, life cycle and mating systems (see Bazin *et al.* (2006) for further details). Additionally, sampling bias may also be responsible for observing different levels of intraspecific genetic diversity and it is already known to decrease the accuracy of PTP (Zhang *et al.*, 2013). To incorporate the potential divergence in intraspecific diversity, we propose the novel multi-rate Poisson Tree Processes (mPTP) model. In contrast to PTP, it fits the branching events of *each* delimited species to a *distinct* exponential distribution. Thereby it can better accommodate the sampling-and population-specific characteristics of a broader range of empirical datasets. In addition, we develop and present a novel, dynamic-programming algorithm for the mPTP model. The implementation is several orders of magnitude faster than the original PTP and yields more accurate delimitations almost instantly on large datasets comprising thousands of taxa. Note that, the original PTP requires days of computation time on a desktop to analyze such datasets. Finally, we provide an *Markov chain Monte Carlo* (MCMC) delimitation sampling approach that allows for inferring delimitation support values.

## 2 METHODS

In this section we introduce the basic notation that is used throughout the manuscript, and provide a detailed description of the mPTP algorithm including the MCMC sampling method.

A binary rooted tree *T* = (*V*, *E*) is a connected acyclic graph where *V* is the set of nodes and *E* the set of branches (or *edges*), such that *E* ⊂ *V* × *V*. Each *inner node u* has a *degree* (number of branches for which *u* is an endpoint) of 3 with the exception of the root node which has degree 2, while *leaves* (or *tips*) have degree 1. We use the notation (*u*, *v*) ∈ *E* to denote an branch with end-points *u*, *v* ∈ *V*, and *ℓ*: *V* × *V* ⟶ ℝ to denote the associated branch length. Finally, we use *T*_*u*_ to denote the subtree rooted at node *u*.

## 2.1 Multi-rate PTP (mPTP) heuristic algorithm

Let *T* = (*V*, *E*) be a binary rooted phylogenetic tree with root node *r*. The optimization problem in the original PTP is to find a connected subgraph *G* = (*V*_*s*_, *E*_*s*_) of *T*, where *V*_*s*_ ⊆ *V*, *E*_*s*_ ⊆ *E*, *r* ∈ *V*_*s*_ such that (a) *G* is a binary tree, and (b) the likelihood of i.i.d branch lengths *E*_*s*_ and *E*_*c*_ = *E* \ *E*_*s*_ fitting two distinct exponential distributions is maximized. Formally, we are interested in maximizing the likelihood

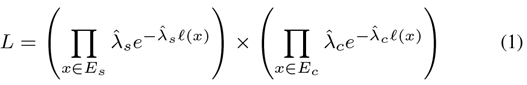

where 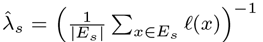 and 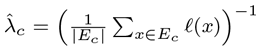 are maximum likelihood (ML) estimates for the rate parameters.

In the multi-rate PTP (mPTP), we are interested in fitting the branch lengths of *each* maximal subtree of *T* (each species delimited in *T*) formed by branches from *E*_*c*_ to a *distinct* exponential distribution. Let *T*_1_ = (*V*_1_, *E*_1_), *T*_2_ = (*V*_2_, *E*_2_), …, *T*_*k*_ = (*V*_*k*_, *E*_*k*_) be the *maximal* subtrees of *T* formed exclusively by branches from *E*_*c*_ such that 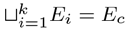 and no pair *i*, *j* exists for which *V*_*i*_ ∩ *V*_*j*_ ≠ ∅. The task in mPTP is to maximize the likelihood

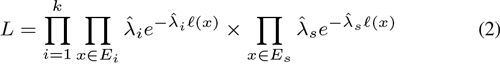

where 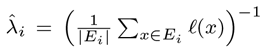 maximal subtree root nodes as *coalescent roots*.

Whether a polynomial-time algorithm exists to solve the two problems remains an open question. We assume that the problem is hard, and thus, we propose a greedy, dynamic-programming (DP) algorithm to solve the problem. The algorithm visits all inner nodes of *T* in a bottom-up postorder traversal, that is, a node is visited only after both of its child nodes have been visited. For each node *u*, the DP computes an array of |*E*_*u*_| + 1 scores, where *E*_*u*_ is the set of branches of subtree *T*_*u*_. Entry 0 ≤ *i* ≤ |*E*_*u*_| assumes that *T*_*u*_ contains *i* between-species branches. For the case *i* = 0, *T*_*u*_ is part of *a* coalescent process, while for *i* = |*E*_*u*_|, *T*_*u*_ is part of *the* single between-species process. Each entry *i* > 0 contains the maximization of Eq. 2, by considering only (i) the branches of subtree *T*_*u*_ and (ii) the set *S* of branches of *T* that have at least one end-point on the path from the root *r* to *u*. For this subset of branches, the between-species process consists of exactly |*S*| + *i* branches, and we only consider the coalescent processes inside *T*_*u*_. We restrict ourselves to the set of branches *S* since it is infeasible to test all possible groupings of branches outside of *T*_*u*_. For any *i* > 0, it holds that the edges from *u* leading to its child nodes *v* and *w* belong to the between-species process, and therefore, *S* is the smallest set of edges which, by definition, must be part of the between-species process. Figure 1 illustrates the algorithm for a particular triplet of nodes *u,v,w*, and marks the set *S* with dashed lines. Entry 0 represents the null-model for *T_u_*, that is, all branches of *T*_*u*_ belong to the coalescent process [or, equivalently, to the speciation process, as the two cases can not be distinguished by the (m)PTP model]. The score (maximization of Eq. 2) for entry *i* ≥ 1 is computed from the information stored in the array entries *j* and *k* of the two child nodes *v* and *w*, by considering all combinations of *j* and *k* such that *i* = *j* + *k* + 2 (plus two accounts for the two outgoing edges of *u*). Not all array entries are necessarily valid. For instance, entry *i* = 1 may be invalid given that node *u* has two out-going branches in the between-species process. Similarly, it is possible that two invalid entries *j* and *k* exist for a given value of *i*. Each entry *i* at node *u* stores the computed score, the sum of between-species branch lengths 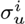 within *T_u_*, the product 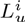 of likelihoods of coalescent processes inside *T*_*u*_, and pointers for storing which entries *j* and *k* were chosen for calculating a specific entry *i*. The first term of Eq. 2 (coalescent process) is the product of 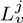 and 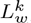, while the second term (speciation process) is computed from the sums 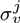 and 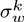 and the sum of branch lengths from *S*. We store the score and child node entry indices for the combination of *j* and *k* that maximize the score. Once the root node of *T* is processed, we have the likelihood values of exactly |*E*| + 1 delimitations, one of which corresponds to the null-model (*i* = 0). We select the entry *i* that minimizes the *Akaike Information Criterion* score corrected for small sample size (AIC_c_;McQuarrie and Tsai (1998)). This way we penalize oversplitting caused by the increasing number of parameters (λ_1_, λ_2_,…, λ_*k*_). A lower AIC_*c*_ value corresponds to a model which better explains the data. Once the best AIC_*c*_ corrected entry *i* at the root has been determined, we perform a backtracking procedure using the stored child node pointers to retrieve the coalescent roots and hence obtain the species delimitation. The average asymptotic run-time complexity of the method is 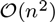 with a worst-case of 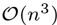, where *n* is the number of nodes in *T* (for the proofs see paragraph 1, Supplement I).

**Fig. 1.**
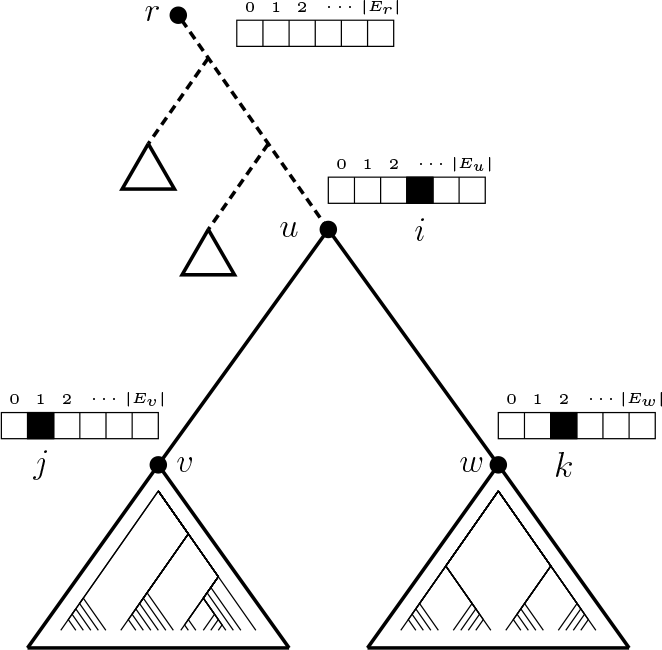
Visual representation of the mPTP dynamic programming algorithm. Each entry *i* at node *u* is computed from information stored at entries *j* and *k* of child nodes *v* and *w*,for all *j* and *k* such that *i* = *j* + *k* + 2. The dashed branches denote the smallest set *S* of branches which, by definition, must be part of the between-species process, irrespective of the resolution of other subtrees outside *T*_*u*_.

## 2.2 MCMC sampling

We deploy an MCMC approach to obtain delimitation support values for each node. In each step, we propose a new delimitation by designating a set of coalescent roots and compute its score.We denote the current delimitation as *θ* and the proposed delimitation, after applying a move to *θ*, as *θ*′. We use the acceptance ratio

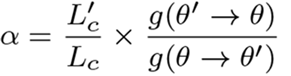

to decide whether to keep the proposed delimitation *θ*′ or not. If *α* ≥ 1, we accept the delimitation, otherwise we accept it with probability *α*. The term 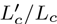 is the ratio of the AIC_*c*_-corrected likelihood values of the *θ*′ over *θ*, and *g*(*x* ⟶ *y*) is the transition probability from one delimitation, *x*, to any other delimitation *y*. Given *θ*, we propose *θ*′ by applying one of two possible moves with equal probability. The first move is to *split* a coalescent clade by placing a randomly selected coalescent root node *u* (and its two out-going branches) in the set of nodes (and branches) corresponding to the speciation process. The second type of move selects a node u whose two child nodes are coalescent roots, *joins* their branch sets into one, and sets *u* as the new coalescent root. The support value for a particular node *u* is the sum of Akaike weights (Akaike, 1978; Burnham and Anderson, 2002) of all MCMC samples for which *u* is part of the speciation process normalized by the sum of all Akaike weights. The method’s run-time is linear to the number of MCMC steps and independent of the tree size. For a sketch of the algorithm see Paragraph 3 of Supplement I.

It is considered good practice that MCMC results are accompanied by a critical convergence assessment. Arguably, a good way of accomplishing this is to compare samples obtained from independent MCMC runs. We use the average standard deviation of delimitation support values (ASDDSV) for quantifying the similarity among such samples. Inspired by the average standard deviation of split frequencies (ASDSF) technique from phylogenetic inference (Ronquist *et al.*, 2012), we calculate the ASDDSV by averaging the standard deviation of per-node delimitation support values across multiple independent MCMC runs that explore the delimitation space. Each such MCMC run starts from a randomly generated delimitation. Similarly to ASDSF, ASDDSV approaches zero as runs converge to the same distribution of delimitations.

ASDDSV is useful for monitoring that chains converge to the same distribution, but does not imply that they have globally converged. It is also desirable to compare the delimitation support values of an MCMC run with the ML estimate. For this, we introduce the *average support value* (ASV), that is,

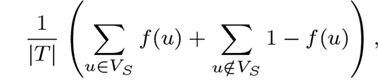

where *V*_*S*_ is the set of the speciation process nodes and *f*(*u*) the support value of a speciation node *u*. The closer ASV is to one, the better the support values agree with the ML delimitation.

## 3 EXPERIMENTAL SETUP

For the evaluation of mPTP we use empirical datasets that comprise a number of varying parameters (substitution rate, geographic range, genetic diversity etc.), which are difficult to reproduce with simulations. We retrieved all datasets from the Barcode of Life Database (BOLD, http://www.boldsystems.org/), the largest barcode library for eukaryotes. For each of the datasets we inferred the putative species with mPTP and four other methods that model speciation on the basis of genetic distance (including the original PTP method), and assess their accuracy with respect to the current taxonomy (available in BOLD). Finally, we avoided comparisons with time-based species delimitation methods (Fujisawa and Barraclough, 2013; Yang and Rannala, 2014; Jones *et al.*, 2015) which are time consuming and heavily dependent on the factorization accuracy of branch lengths into time and evolutionary rate.

**Table 1.**
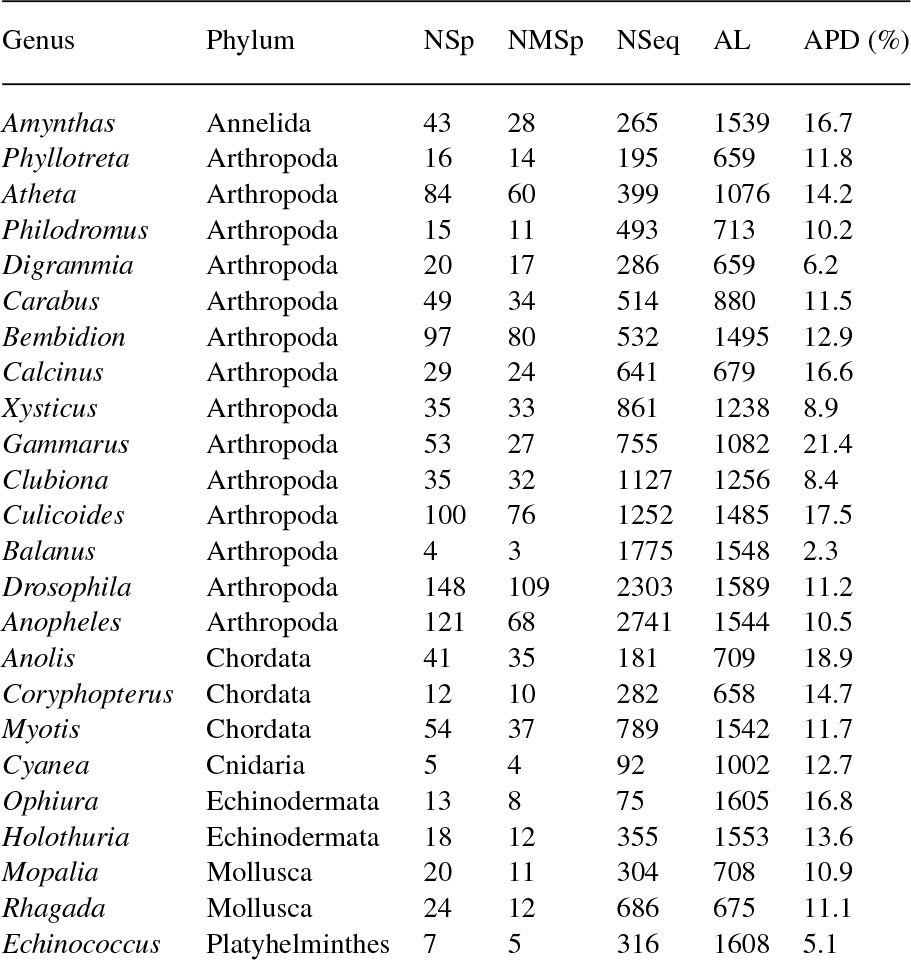
Main characteristics of empirical test datasets. NSp: Number of species, NMSp: Number of Monophyletic species, NSeq: Number of Sequences, AL: Alignment Length, APD: Average P-Distance.

## 3.1 Empirical Datasets

We retrieved 24 empirical genus level datasets (Table 1) that cover six of the most species rich animal Phyla (Arthropoda, Annelida, Chordata, Echinodermata, Platyhelminthes, Cnidaria) in BOLD. This allows to evaluate the efficiency of mPTP given a wide range of organisms with diverse biological traits and evolutionary histories. The majority of datasets (14 out of 24 datasets) belong to the Arthropoda which is the most species-rich animal group, and the most common group PTP has been applied to so far (in 61 out of 110 citations). Within Arthropoda, we mainly focus on insects (8 out 14 datasets), which represents over 90% of all animals. The rest of the empirical datasets spans the remaining five phyla.

All datasets comprise the 5’ end of the common animal barcoding gene “Cytochrome Oxidase subunit I” (COI-5P) (Hebert *et al.*, 2003). The number of taxonomic species in the 24 datasets ranged from four (*Balanus*) to 148 (*Drosophila*), while the number of sequences from 75 (*Ophiura*) to 2741 (*Anopheles*). The number of sequences per species ranged between two and the extreme case of 1763 for *Balanus glandula*, reflecting situations where some species might be readily and others rarely available. Finally, the geographical distribution of the datasets also varied from the global scale (e.g., *Anopheles, Drosophila*) to the local scale (e.g., the *Anolis* samples originate from the islands and surrounding shores of the Caribbean sea).

## 3.2 Putative Species Delimitation

The sequence files obtained from the BOLD database were preprocessed to remove three factors that interfere with the assessment of the delimitation methods. First, the efficiency of each method is measured by comparing the delimited to the taxonomic species (see paragraph 3.3). Therefore, it was necessary to remove sequences of ambiguous taxonomic assignment. Second, we removed duplicate sequences, using the “-f c” option of RAxML version 8.1.17 (Stamatakis, 2014), to avoid unnecessary computations. Finally, we removed *singleton species* (i.e., species represented by a single sequence) to eliminate the impact of random exact matches (of delimited to taxonomic species) in methods that tend to oversplit. Additionally, singletons also interfere with the efficiency of delimitation methods (Puillandre *et al.*, 2012).

For each dataset, we inferred putative species using the two PTP models and we compared the results to three popular and well-established distance-based methods; ABGD (Puillandre *et al.*, 2012), Uclust (Edgar, 2010) and Crop (Hao *et al.*, 2011). The input format of the data and parameters differ among the five methods. The two PTP versions require a rooted phylogenetic tree. Therefore, we used Mafft v7.123 (Katoh and Standley, 2013) to align the sequences of each dataset, and subsequently RAxML under the default algorithm and the GTR+Гmodel, to infer an ML tree. To root each tree, we chose outgroup taxa based on previously published phylogenies. For the genera without such prior phylogenetic knowledge, we selected a representative species of a genus belonging to the same family or tribe given the taxonomy in BOLD (NCBI Accession Numbers of the ingroup and outgroup sequences are provided in Supplement II). Moreover, we implemented a method that identifies and ignores branches that result from identical sequences, during the delimitation process (for more details see paragraph 2 and Figures 1 and 2, Supplement I).

For mPTP, we further assessed the confidence of the ML solution using MCMC sampling. For each dataset, we executed ten MCMC runs of 2 × 10^7^ steps, each starting from an initial random delimitation. We estimated the convergence of the independent runs by calculating the ASDDSV. To obtain an overall support for the ML estimate, we computed the mean ASV over all ten independent runs.

For each of the remaining three methods (ABGD, Uclust and Crop), we optimized a set of performance-critical parameters and chose the delimitation that recovered the highest number of taxonomic species. ABGD requires two parameters: (i) the prior limit to intraspecific diversity (*P*) and (ii) a proxy for the minimum barcoding gap among the inter-and intraspecific genetic distances (*X*). We ran ABGD with the aligned sequences and 400 parameter combinations. For *P* we sampled 100 values from the range 〈0.001, 0.1〉 and for *X* we used the four values 0.05, 0.1, 0.15 and 0.2. The only critical parameter for Uclust is the fractional identity threshold (*id*) which defines the minimum similarity for the sequences of each cluster. We performed 50 consecutive runs, increasing the *id* value by 0.01 starting with 0.5. Finally, for Crop we tried four combinations of the *l* and *u* parameters that correspond to similarity thresholds of 1%, 2%, 3%, and 5% as suggested by the authors (http://code.google.com/p/crop-tingchenlab/). Another critical parameter for Crop is *z*, which specifies the maximum number of sequences to consider after the so-called initial “split and merge”. We set *z* to 100 which is considered as a reasonable value for full-length barcoding genes.

## 3.3 Comparison of delimitation methods

The evaluation of the five methods is based on three measures. First, the percentage of recovered taxonomic species (RTS), that is, the percentage of delimited species that match the “true species”. We consider the taxonomic species retrieved from BOLD to be the “true species”, and we deem the performance of the algorithm better when the number of matches to the taxonomic species is higher. Since we do not have the expertise to evaluate the taxonomy of each dataset, we could not assess the accuracy of each individual delimitation. However, by assuming that, the closer a delimitation is to the current taxonomy, the higher the probability to correspond to the real species, we gain some insight as to the relative accuracy among the different methods. The second measure is the F-score, also known as F-measure or F1-score (Rijsbergen, 1979), that is, the harmonic mean of *precision* and *recall* measures. In species delimitation, *precision* denotes the fraction of clustered sequences belonging to a single taxonomic species. The *recall,* describes the fraction of sequences of a species that are clustered together. The F-Score improves when decreasing (i) the number of species which are split into more than one groups and (ii) the number of taxonomic species lumped together into one group. The F-score ranges from 0 to 1, where 1 indicates a perfect agreement among two delimitations. Finally, the third measure is simply the number of delimited species.

**Fig. 2.**
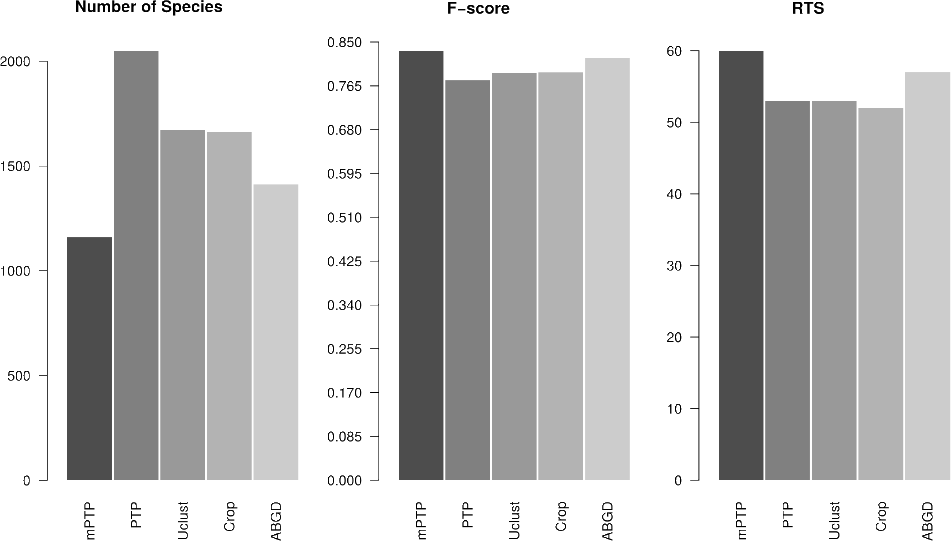
Average performance over all datasets of the five delimitation methods (mPTP, PTP, Uclust, Crop, and ABGD) for the A) number of species, B) F-scores and C) number of RTS

The accuracy of either PTP model in recovering the true species depends on whether these taxa form monophyletic clades. Thus, for PTP and mPTP, we also compare their performance by applying the above measures for monophyletic taxonomic species only. Finally, we also quantify how the percentage of monophyletic species correlates to the percentage of RTS for the five delimitation methods we test.

## 4 RESULTS

We analyzed a total of 17,219 COI sequences. The alignment length ranged from 658 bp to 1620 bp, while the proportion of variable nucleotides, measured by the P-Distance (MEGA v5.2; Tamura *et al.*, 2011) averaged over the sequences of each dataset, varied from 2.3% (*Balanus*) to 21.4% (*Gammarus*). Table 1 presents the fraction of monophyletic taxonomic species in the phylogenies for each dataset. The fraction ranges from only 51% in *Gammarus*, the most variable of the datasets, to 94% (*Xysticus*). Out of the 400 combinations of parameters tested for ABGD, two recovered the highest number of species, with *P* = 0.010235 and *X* = 0.1 or 0.05. For Uclust, the delimitation that maximized the number of RTS was with the threshold value of *id* = 0.97. Finally, the best scoring parameter combination for Crop (*l* = 1.0 and *u* = 1.5) corresponds to the 3% similarity threshold.

Table 2, presents the efficiency of the five delimitation methods for five of the datasets using three measures: percentage of RTS, F-score and number of species. A complete list of results for all 24 datasets is given in paragraph 4.1 in Supplement I. Overall, mPTP and ABGD scored best in terms of RTS percentage (59% and 57% on average, respectively) and F-scores (0.828 and 0.819, respectively) compared to the other three methods (PTP: RTS = 53%, F-score = 0.776, Uclust: RTS = 53%, F-score = 0.79, Crop: RTS = 52%, F-score = 0.791) (Figure 2). The striking difference between the five methods is in the number of delimited species. The novel mPTP method delimited a total of 1190 species which is the closest to the total number of taxonomic species (1041). In contrast, PTP inferred 2048 species which is almost twice this number. The other three methods yielded more conservative species numbers compared to PTP (Uclust: 1671, Crop: 1663, ABGD: 1412) that are, however, still notably higher than the mPTP estimates (Figure 2).

When considering only monophyletic species, as one might expect, the recovery percentage increases substantially for both PTP models (82% and 72% on average for mPTP and PTP, respectively), indicating that polyphyly [we use the term polyphyly in referring to both paraphyly and polyphyly *sensu* Funk and Omland (2003)] is a major contributing factor when taxonomic species are not recovered (Figure 3 in Supplement I). In particular, the RTS percentage of either PTP model is highly correlated with the percentage of monophyletic species in a dataset (Figure 3). The Pearson coefficient indicates that the correlation is stronger among the species recovered with mPTP (*r*_*m*_*PTP* = 0.75, p-value = 2.642e-05) than with PTP (*r*_*PTP*_ = 0.62, p-value = 0.001218). The correlation is also positive for the other three methods (*r*_*ABGD*_ = 0.63 / p-value = 0.0009327, *r*_*Uclust*_ = 0.5 / p-value = 0.01294, *r*_*Crop*_ = 0.63 / p-value = 0.0009421) and comparable to PTP but smaller than for mPTP. Similarly, the slope of the regression line was greater for mPTP than for the other methods, indicating a steeper linear relationship between the two variables (Figure 3).

**Table 2.** Percentage of RTS, F-scores and number of delimited species for the five delimitation methods (mPTP,PTP,Uclust, Crop and ABGD) for five of the empirical datasets.

|  | mPTP | PTP | Usearch | Crop | ABGD |
| --- | --- | --- | --- | --- | --- |
| **RTS (%)** |  |  |  |  |  |
| *Amynthas* | 51 | 40 | 44 | 40 | 44 |
| *Anopheles* | 41 | 39 | 30 | 31 | 40 |
| *Atheta* | 64 | 60 | 57 | 55 | 60 |
| *Drosophila* | 47 | 51 | 41 | 34 | 53 |
| *Philodromus* | 60 | 27 | 47 | 53 | 47 |
| **F-score** |  |  |  |  |  |
| *Amynthas* | 0.784 | 0.638 | 0.649 | 0.674 | 0.673 |
| *Anopheles* | 0.787 | 0.704 | 0.730 | 0.681 | 0.730 |
| *Atheta* | 0.839 | 0.844 | 0.836 | 0.832 | 0.850 |
| *Drosophila* | 0.728 | 0.559 | 0.747 | 0.729 | 0.765 |
| *Philodromus* | 0.882 | 0.717 | 0.812 | 0.852 | 0.828 |
| **Number of species** |  |  |  |  |  |
| *Amynthas* | 64 | 104 | 95 | 91 | 90 |
| *Anopheles* | 126 | 218 | 193 | 154 | 118 |
| *Atheta* | 85 | 100 | 95 | 95 | 91 |
| *Drosophila* | 139 | 444 | 183 | 217 | 157 |
| *Philodromus* | 21 | 38 | 26 | 20 | 24 |

Regarding the confidence of the mPTP delimitations, all independent MCMC runs appear to converge with an ASDDSV below 0.01 for all datasets, except for *Drosophila* (one of the largest datasets), for which the ASDDSV was 0.048, but still in the acceptable range for assuming convergence (Ronquist *et al.*, 2012). The ASV with respect to the ML delimitation was very high for all datasets, ranging from 72% (*Myotis*) to 99.8% (*Phyllotreta*), indicating that the data support well the ML solution (Table 3 in Supplement I). The accumulated running time for all ten independent runs (executed sequentially on an Intel Core i7-4500U CPU @ 1.80GHz) was less than 50 seconds on average across all datasets, which corresponds to ~ 5 seconds per run. For a thorough run-time comparison between PTP and mPTP (including the ML method) please refer to paragraph 4.2 in Supplement I.

**Fig. 3.**
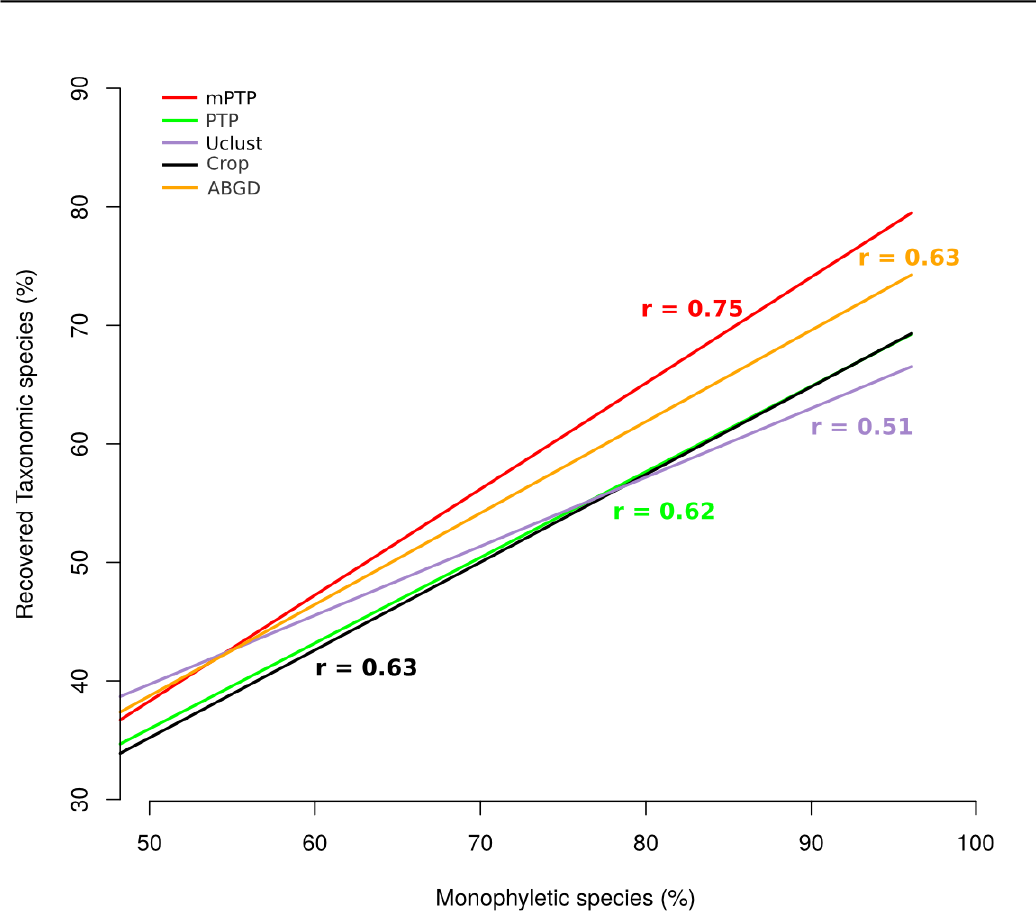
For each method, we fit a regression line to the points of correspondence of the percentage of monophyletic species (x-axis) to the percentage of RTS (y-axis). The Pearson coefficient (*r*) is given for each correlation in the corresponding color.

## 5 DISCUSSION

Molecular species delimitation has caused mixed reactions within the scientific community, from those highly enthusiastic about its potential to accelerate biodiversity cataloging (Blaxter, 2003; Blaxter and Floyd, 2003) to those very critical about its role in shaping modern systematics (Bauer *et al.*, 2010; Will *et al.*, 2005). The main argument between the two conflicting sides is whether molecular delimitation on its own is sufficient to justify taxonomic rearrangements (Will *et al.*, 2005). Integrative taxonomy alleviates this conflict as it, by definition, requires multiple levels of evidence taking into account various biodiversity characteristics of an organism to accept potential taxonomic changes. Within this framework, and in line with the independently evolving species concept (De Queiroz, 2007) and the phylogenetic species concept, molecular species delimitation using DNA-barcoding serves as an excellent tool in modern taxonomy (Tautz *et al.*, 2003; Vogler and Monaghan, 2007). It can be easily applied to a large range of organisms regardless of their life stage, gender or prior taxonomic knowledge, by a broad range of researchers. In addition, barcoding genes are easily amplified from small tissue samples even from poorly preserved historical samples (Austin and Melville, 2006). Furthermore, such an approach might represent the only feasible approach in biodiversity surveys, as they often comprise a large number of species, many of which might be unknown or not easily accessible. Hence, collecting ecological or morphological traits is often simply not feasible. For historical samples or samples from inaccessible areas (e.g., deep seas, deserts) barcoding methods are equally important since the collection of life history traits might be similarly challenging. This and previous studies (Monaghan *et al.*, 2009; Esselstyn *et al.*, 2012; Tang *et al.*, 2014; Ratnasingham and Hebert, 2013) suggest that single-locus barcoding methods provide meaningful clusters, close to taxonomically acknowledged species. This makes them useful for approximate species estimation studies or when more thorough systematic research is practically impossible.

### 5.1 (m)PTP

The molecular systematics of the example taxa, vary from well-studied [e.g., *Anopheles* (Harbach and Kitching, 2016), *Drosophila* (Bächli, 2016)] to scarcely studied (e.g., *Xysticus*, *Clubiona*). They further differ in the number of species, number of sequences per species, geographic ranges, and nucleotide divergences. Despite these differences, mPTP outperformed PTP and yields substantially smaller putative species numbers as well as delimitations that are closer to the current taxonomy. The assumption of mPTP that per-species branch lengths can be fit to distinct exponential distributions increases flexibility and adjustability to more realistic datasets (e.g., including variable intraspecific diversity). The divergence of intraspecific diversity patterns may either be due to population traits and processes (Bazin *et al.*, 2006) or the uneven sampling of the species (e.g., a highly sampled species from a single location compared to a species represented by one sample from multiple locations). The latter is known to decrease PTP accuracy (Zhang *et al.*, 2013).

The accuracy of (m)PTP strongly correlates with the proportion of monophyletic species in the underlying phylogeny. Paraphyletic species will either be delimited into smaller groups or delimited with other, nested species. Therefore, the recovery rate of taxonomic species is significantly higher when only considering the monophyletic species in each dataset. Among the 24 datasets, the number of monophyletic species ranged from 51% to 94%. Thus, the lack of monophyly is the primary contributing factor for not recovering taxonomic species with either PTP model. The average observed monophyly in our arthropod (76.4%) and the remaining invertebrate (64.7%) datasets corresponds well to available estimates for the same groups (73.5% and 61.4%, respectively) based on a large number of empirical studies (Funk and Omland, 2003). The same study reports that the most common reasons for polyphyly are: inaccurate taxonomy, incomplete lineage sorting, and retrogressive hybridization (Funk and Omland, 2003; McKay and Zink, 2010). All three effects could apply to the selected datasets. Nevertheless, only the latter two are relevant to the efficiency of the algorithm *per se*, while the accuracy of the taxonomy is only relevant when it is used as a reference measure. Polyphyletic species affect both PTP versions in the same way. The reason for the improved delimitation accuracy of monophyletic species of mPTP over PTP ie because it can accommodate different degrees of intraspecific genetic diversity within a phylogeny.

### 5.1.1 MCMC Sampling

The drawback of an ML approach is that it only provides a point estimate and no information on model uncertainty. The confidence about a given solution has a substantial impact in drawing reasonable conclusions. Therefore, we provide an MCMC method for assessing the plausibility of the ML solution. In phylogenies comprising hundreds of taxa, it is hard to obtain an overall support for a particular delimitation hypothesis by visual inspection of the tree. To alleviate this problem, at the end of an MCMC run we calculate the ASV for the ML solution. In our experiments, the ML delimitations were highly supported by the ASVs for all datasets, pointing towards a unimodal likelihood surface for our model (Table 3 in Supplement I). Low ASVs may be interpreted as low confidence for the given delimitation scheme, either because another (multi-modal likelihood surface) or no delimitation scheme (flat likelihood surface) is well supported. This also indicates a poor fit of the data to the model. The execution times of the MCMC sampling are almost negligible, regardless of tree size. Therefore, we can thoroughly sample the delimitation space even for phylogenies comprising thousands of taxa. For the large phylogenies of our study (i.e., *Drosophila*, *Anopheles*), the PTP implementation required over 30 hours for the ML optimization alone, and would require days or even months for the estimation of support values. Instead, mPTP required less than a minute for both the ML optimization and the support value estimation.

### 5.2 Distance-based Methods

Distance-based methods are easy to apply to large datasets as they need minimal preprocessing effort and computational time. Their major weakness is that they require either a threshold value or a combination of parameters associated with the threshold value, the sampling effort, or the search strategy. Empirical data show that certain similarity cut-offs (2–3%) correspond well to the species boundaries of several taxonomic groups (Hebert *et al.*, 2003; Smith *et al.*, 2005; Hebert *et al.*, 2004); however, these values are often far from optimal (Lin *et al.*, 2015). Selecting these parameters is not intuitive and they may only be evaluated *a posteriori* based on the expectations of the researcher. Furthermore, the empirical knowledge for threshold settings is tightly associated with barcoding genes, and it may not be as useful for other marker genes. Here, we optimized the parameters for three of the most popular distance methods (ABGD, Crop and Uclust) based on the RTS percentage. Despite our substantial effort to use optimal parameters for the given data, our results show that, on average, mPTP performs better with respect to F-scores and RTS percentage. At the same time, it also delimited notably fewer species. ABGD accuracy was closest to mPTP, while PTP, CROP, and Uclust performed notably worse. Besides accuracy, the greatest advantage of mPTP is that it consistently yields more accurate results without requiring the user to optimize any parameters/thresholds. Finally, in contrast to (m)PTP, distance methods ignore evolutionary relationships. Hence, there is no direct relation between monophyletic species and delimitation accuracy in similarity based tools. However, polyphyly often reflects recent speciation while reciprocal monophyly indicates that significant time since speciation has passed. Consequently, the barcoding gap should be less pronounced in datasets of recently diverged species. This justifies that the RTS fraction improves with the number of monophyletic species.

### 6 CONCLUSIONS

We presented mPTP, a novel approach for single-locus delimitation that consistently provides faster and more accurate species estimates than PTP and other popular delimitation methods. As PTP, mPTP is mainly designed for analyzing barcoding loci, but can potentially also be applied to entire organelle phylogenies (e.g. mitochondria, Qin *et al.*, 2015). In contrast to methods based on sequence similarity, mPTP does not require any similarity threshold or other user-defined parameter as input. The limitations of mPTP are associated with processes that can not be detected neither in single-gene phylogenies (incomplete lineage sorting, hybridization) nor in recent speciation events. The novel dynamic programming delimitation algorithm reduces computation time to a minimum and allows for almost instantaneous species delimitation on phylogenies with thousands of taxa. The MCMC sampling provides support values for the delimited species based on millions (or even billions) of MCMC generations in just a few seconds on a modern desktop. The mPTP tool is available both, as a standalone package, and as a web service.

## ACKNOWLEDGMENT

The authors gratefully acknowledge the support of the Klaus Tschira Foundation. PP was funded by FP7-PE0PLE-2013-IEF EVOGREN (625057).

